# The AKT Forkhead box O transcription factor axis regulates human cytomegalovirus replication

**DOI:** 10.1101/2022.04.14.488435

**Authors:** Hongbo Zhang, Anthony J. Domma, Felicia D. Goodrum, Nathaniel J. Moorman, Jeremy P. Kamil

## Abstract

The serine/threonine protein kinase AKT is a critical mediator of growth factor signaling that broadly impacts mRNA translation, metabolism, programmed cell death and stress responses. Human cytomegalovirus (HCMV), a large dsDNA virus in the herpesvirus family, dramatically remodels host cell signaling pathways during lytic infection. Here, we show that AKT accumulates in an inactive form during HCMV infection, as indicated by hypo-phosphorylation at Thr308 and Ser473, as well as nuclear localization of FOXO3a. Moreover, we find that expression of constitutively active myristoylated AKT (myr-AKT) causes a 1-log defect in viral replication, accompanied by defects in viral DNA synthesis and viral late gene expression. These findings suggest that HCMV shutoff of AKT is not merely incidental to viral factors such as pUL38 that maintain high levels of mTORC1 activity independently of extracellular growth factor signals. Short interfering RNA knockdown of FOXO3a, but not FOXO1, phenocopied the replication defects seen during expression of myr-AKT, corroborating a role for FOXO3a during HCMV infection. Accordingly, a chimeric FOXO3a-estrogen receptor fusion protein, whose nuclear localization is regulated by the small molecule ligand 4-hydroxytamoxifen instead of AKT activity, rescues virtually all the replication defects induced by myr-AKT expression. Taken together, our results argue that HCMV dampens AKT signaling to promote nuclear localization of FOXO transcription factors, which are required for efficient viral replication.

**IMPORTANCE:** Evidence from a diverse herpesvirus infection models suggests that the PI3K / AKT signaling pathway suppresses reactivation from latency, and that inactivation of the pathway stimulates viral lytic replication. Here, we show that AKT accumulates in an inactive state during HCMV infection of lytically permissive cells while the presence of constitutive AKT activity causes substantial viral replication defects. Although AKT phosphorylates a diverse array of cellular substrates, we identify an important role for the Forkhead box class O transcription factors. Our findings show that when FoxO3a nuclear localization is decoupled from its negative regulation by AKT, the viral replication defects observed in the presence of constitutively active AKT are reversed. Collectively, our results reveal that HCMV inactivates AKT to promote the nuclear localization of FOXO transcription factors, which strongly implies that FOXOs play critical roles in transactivating cellular and/or viral genes during infection.

## INTRODUCTION

The serine/threonine protein kinase AKT regulates a broad range of cellular activities in response to signaling by receptor tyrosine kinases and G-protein coupled receptors [reviewed in (1)]. For instance, AKT enhances mRNA translation by phosphorylating PRAS40 and TSC2, which otherwise constrain mTORC1 activity (2, 3). Meanwhile, AKT promotes cell survival by phosphorylating the pro-apoptotic Bcl-2 family protein BAD (4). AKT directly phosphorylates various enzymes and proteins that regulate metabolism. For example, AKT phosphorylation of HK2 stimulates conversion of glucose to glucose-6-phosphate while its phosphorylation of TXNIP inhibits endocytosis of glucose transporters from the plasma membrane (5). Although the positive influence of AKT on mRNA translation and inhibition of apoptosis might be expected to promote viral replication, a number of cytolytic viruses, such as measles virus and vesicular stomatitis virus inactivate AKT during infection (6–8). Precisely how viruses benefit from shutting off AKT is unknown. However, it has been proposed that viruses might inactivate AKT to inhibit the translation of transcripts encoding interferon stimulated genes (9).

AKT is activated at very early times during human cytomegalovirus (HCMV) entry into monocytes (10), and initial studies in lytically infected fibroblasts suggested that AKT remains active during lytic infection (11, 12). Nonetheless, more recent findings show that AKT is, in fact, inactive during infection in permissive human fibroblasts (13, 14), even though downstream processes that ordinarily require AKT remain active. For example, HCMV expresses a protein called pUL38 that inactivates TSC2, which provides a mechanism to maintain mTORC1-mediated stimulation of mRNA translation without AKT (15). Yet, whether the virus benefits from maintaining AKT in an inactive state has remained unclear. One class of AKT substrates that may shed light on this unresolved matter is the Forkhead box O family transcription factors (FOXOs).

FOXOs coordinate transcriptional responses to metabolic and oxidative stress, and strongly impact the expression of genes involved in immunity, apoptosis, cell differentiation, and cell cycle progression (16). AKT phosphorylates FOXO3a at Thr32, Ser253 and Ser315; phosphorylation at these sites recruits the binding of 14-3-3 proteins, which exclude FOXOs from nucleus (17, 18).

During growth factor withdrawal or oxidative stress, AKT activity ceases and FOXOs accumulate in the nucleus to transactivate stress response genes. We recently reported that FOXO3a plays roles in transactivating major immediate early (MIE) gene expression during HCMV reactivation from latency in myeloid cells (19, 20). Here, we investigate the significance of AKT regulation to lytic HCMV replication, specifically addressing its regulation of FOXO3a. We find that AKT activity is sharply downregulated during HCMV infection and that expression of a constitutively active AKT, myrAKT, represses HCMV replication with corresponding reductions in viral gene expression and viral DNA synthesis. We further show that FOXO3a, but not FOXO1, is the key substrate of AKT important for HCMV replication. FOXO3a is inactivated by AKT and its forced localization to the nucleus overcomes the defect in replication imposed by constitutive active AKT. These findings establish FOXO3a as a crucial target underlying HCMV-control of AKT and FOXO3a as a pivotal host transcription factor important to HCMV replication.

## RESULTS

### Akt is inactivated during HCMV infection of permissive cells

Akt, also known as protein kinase B, is negatively regulated during the productive (or ‘lytic’) HCMV replication cycle, and its kinase activity is suppressive to replication (13, 14). To further explore how HCMV regulates Akt activity, we infected telomerase-immortalized human fibroblasts with HCMV strain TB40/E at an MOI of 2 TCID50/cell and used phospho-specific antibodies to monitor phosphorylation of Akt at two sites important for its kinase activity, Ser473 and Thr308 (21–23) (**Fig 1A-1B**). We found that Akt phosphorylation at Ser473 markedly decreased by 3 hours post infection (hpi) and by 12 hpi for Thr308. Measurements of signal intensity from far-red fluorescent dye conjugated secondary antibodies allow us to estimate that Akt phosphorylation at Ser473 decreased to 53% of its ‘time zero’ value by 3 hpi, was down to 34% by 6 hpi, to 21% by 12 hpi, by 24 hpi and 48 hpi decreased to 3.3% and 1.9% of its initial intensity, respectively (**Fig 1B, SI Fig S1**). Phosphorylation signal for Thr308 decreased to approximately half (42.5%) of its initial value by 12 hpi, and by 24 hpi and 48 hpi had decreased to 11.4% and 3.4% of its initial value, respectively, while total Akt levels remained roughly constant. Phosphorylation of the proline-rich Akt substrate of 40 kD (PRAS40) at Thr246, a canonical indicator of Akt activity (24), decreased in lockstep with Akt phosphorylation status, while the total abundance of PRAS40 held constant. This suggests that decreased Akt phosphorylation at Thr308 and Ser473 coincided with a decrease in Akt-mediated phosphorylation of one of its cellular substrates (**Fig 1B**).

**Figure 1.**
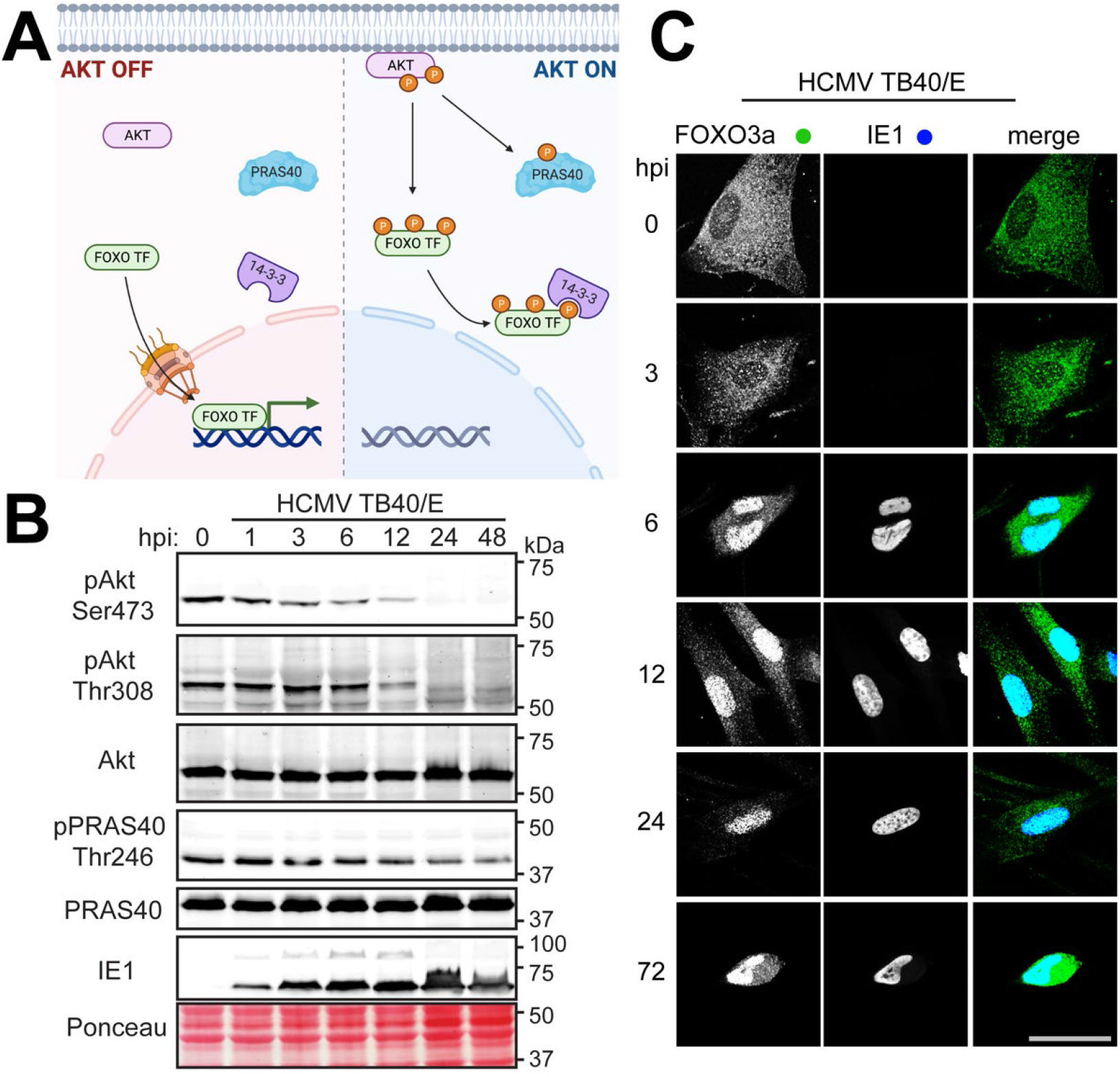
AKT is inactivated during HCMV infection. (**A**) Schematic denoting the influence of Akt activity on two key substrates, PRAS40 and FOXO transcription factors (FOXO TF). Left panel: when Akt is inactive, FOXO TF are able to enter the nucleus. Right panel: when Akt is recruited to membranes, such as during PI3K signaling, Akt is phosphorylated at activating residues Thr308 and Ser473. Once activated Akt in turn phosphorylates many downstream substrates, including PRAS40 and FOXO TF. Akt phosphorylation of FOXO TF recruits binding of 14-3-3 proteins, which sequester FOXO TF in the cytoplasm. (**B**) Fibroblasts were infected with HCMV strain TB40/E at MOI = 2. Lysates were harvested at the indicated times post infection (h post infection, hpi) and analyzed by Western blot for proteins immunoreactive to antibodies specific for pAkt; pAkt Ser473 and pAkt Thr308: Akt phosphorylated at Thr308 or Ser473, respectively; Akt: total Akt; PRAS40 Thr246: PRAS phosphorylated at Thr246, PRAS40: total PRAS40, IE1: the 72-kD HCMV immediate-early nuclear antigen, IE1. A section of nitrocellulose membrane was also subjected to Ponceau S staining (Ponceau) to assess total protein loading across lanes. (**C**) Fibroblasts were infected with HCMV strain TB40/E at MOI = 1. Cells were fixed in 4% PFA at the indicated times post-infection and stained with antibodies specific for FOXO3a (green) or HCMV IE1 antigen (blue). Scale bar: 25 µm.

We also monitored the Akt substrate, forkhead box O transcription factor, FOXO3a, by indirect immunofluorescence staining. Akt negatively regulates the nuclear localization of FOXO family transcription factors (17, 25, 26). Therefore, we used accumulation of FOXO3a in the nucleus as an orthogonal indicator for inactivation of Akt (**Fig 1A**). Although FOXO3a signal was detected predominantly within the cytoplasm at 1 hpi and 3 hpi, the protein localized strongly to the nucleus by 6 hpi and remained there throughout the remainder of the 72 h infection time course (**Fig 1C**). From experiments, we concluded that HCMV infection of permissive human fibroblasts is associated with accumulation of Akt in an inactive form, in agreement with previous observations (13, 14).

### Constitutive AKT activity inhibits HCMV replication

To address whether shut-off of Akt kinase activity is required for efficient HCMV replication, we made use of a constitutively active form of Akt, myr-Akt, which carries a retroviral myristoylation signal at its N-terminus such that localization to membranes, the rate-limiting step in Akt activation, is no longer regulated by PI3K. Making use of permissive human fibroblast cells stably transduced with “tet-on” expression cassettes that enable myr-Akt expression to be induced with doxycycline (dox), we expressed either myristoylated Akt1 (myr-Akt) or a ‘kinase-dead’ K179M myr-Akt control for 24 h prior to infection at MOI = 1 with either of two different HCMV strains, AD169rv and TB40/E. This system allowed us to ask whether the presence of constitutive Akt activity would impact the production of infectious particles over a six day time course. As expected, HCMV infection of cells expressing kinase dead K179M myr-AKT showed robust nuclear localization of FOXO3a at 72 hpi, while cells expressing myr-AKT failed to show FOXO3a localization to the nucleus, consistent with the known role of Akt kinase activity in suppressing nuclear localization of FOXO transcription factors (**Fig 2C-E, SI Fig S2**). However, the two HCMV strains each exhibited a roughly 10-fold replication defect in cells expressing myr-AKT, but not the kinase-dead control (**Fig 2A-B**). From these results, we concluded that the presence of myr-Akt during HCMV infection caused a substantial viral replication defect.

**Figure 2.**
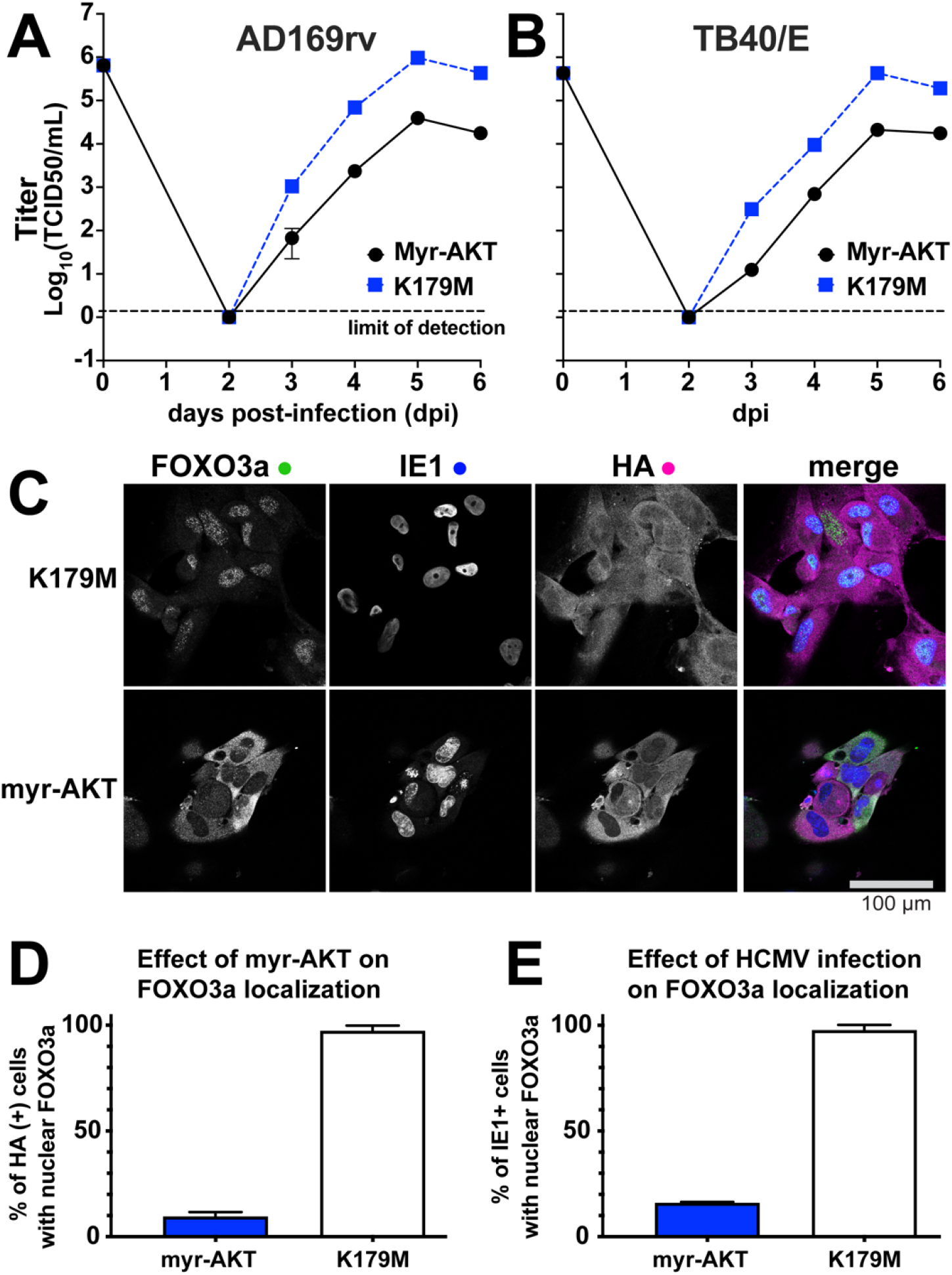
Constitutive AKT activity causes a viral replication defect. (A-B) Fibroblasts carrying doxycycline inducible (“tet-on”) expression cassettes for either myristoylated AKT1 (myr-AKT) or a kinase dead control (K179M) were induced for 24 h using 100 ng/mL doxycycline hyclate and subsequently infected at MOI =1 with either HCMV strain AD169rv (A) or TB40/E (B). Supernatants were harvested at the indicated times post infection (days post infection, dpi) and measured for the infectious titer by TCID50 assay. (C) Representative confocal microscope images of formalin fixed cells at 24 h post infection, see Supp Fig 1 for full time course. (D-E) At least 30 cells per condition were scored at 72 hpi for the effect of myr-AKT or myr-AKT-K179M expression (scored by HA epitope positivity in indirect immunofluorescent staining, the myr-AKT transgene carries an HA tag), or for the effect on HCMV infection (scored by IE1 antigen positive cells) on nuclear localization of endogenous FOXO3a, as indicated on the Y-axis.

### Constitutively active AKT causes defects in viral gene expression and viral DNA synthesis

To better understand the nature of viral replication defect that occurs during expression of constitutively active AKT, we next analyzed a selected set of viral RNA transcripts and proteins during HCMV strain TB40/E infection of fibroblasts (MOI 1) expressing myr-Akt or a kinase dead control. Reverse-transcriptase quantitative PCR (RT-qPCR) results indicated that *UL123* (IE1) mRNA levels accumulated indistinguishably in myr-Akt and ‘kinase dead’ myr-Akt control settings (**Fig. 3A**).

**Figure 3.**
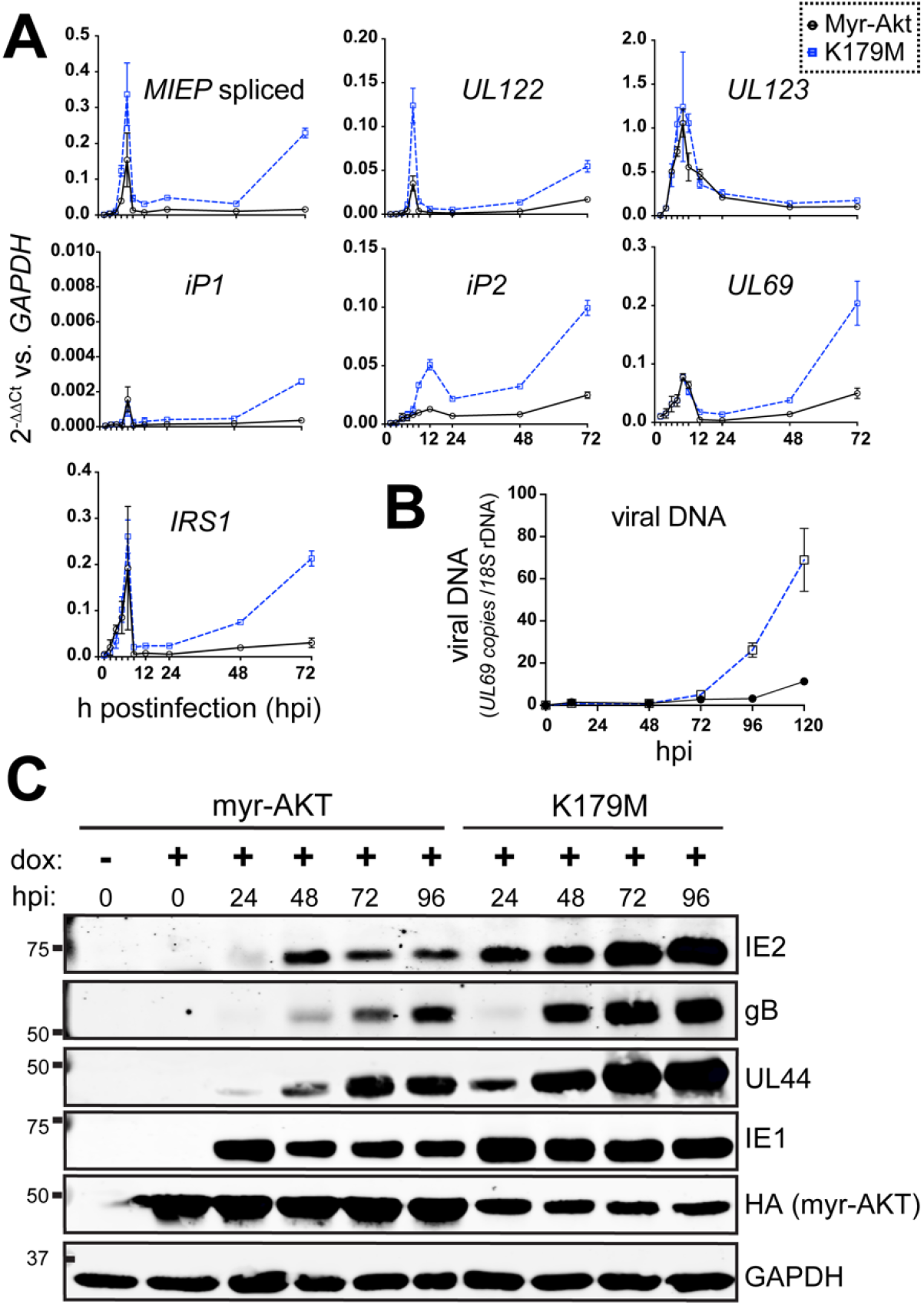
Constitutive AKT activity causes defects in viral gene expression and viral DNA synthesis. (**A**) Fibroblasts were induced for expression of either myr-AKT or myr-AKT-K179M (kinase dead) for 24 h and then were infected with HCMV strain TB40/E at MOI = 1. and total RNA were isolated from either myr-AKT or K179M settings at 2, 4, 6, 8, 10, 12, 16, 24, 48, and 72 hpi, including an on-column DNAse I digestion step. RNA samples were reverse-transcribed into cDNA and assayed by qPCR for the abundance of the indicated viral transcripts relative to levels of cellular *GAPDH* mRNA. (**B**) Total DNA was isolated at the indicated times post infection (hpi) from cells infected exactly as described for panels A and B, and copies of viral DNA were enumerated using quantitative PCR for the HCMV *UL69* gene normalized to copies of cellular 18S rDNA loci. (**C**) Infections were set up as in panel A. Cell lysates collected at the indicated times post-infection were assayed by Western blot for the expression of the indicated viral proteins, as well as for the expression of myr-AKT (HA tag) or GAPDH, as a loading control.

However, *UL122* transcripts, which encode the 86-kD viral MIE transactivator protein IE2, accumulated to decreased levels during infection of myr-Akt expressing cells, but not during infection of comparator cells expressing a ‘kinase dead’ myr-Akt (K179M). A similar pattern was seen for total levels of spliced transcripts originating from the major immediate early promoter (MIEP). Notably, our previous results from hematopoietic cell infection models indicate that FOXO3a transactivates promoters within intron A of the canonical MIE locus, contributing to the accumulation of *IE2 (UL122)* mRNA and protein (19, 20); and these promoters, iP1 and iP2, were also previously found to contribute to IE2 expression during lytic infection of fibroblasts (27). The intronic promoters, particularly iP2, showed decreased activity in the presence of myr-Akt (**Fig. 3A**). Likewise, mRNAs for two other IE genes, *UL69* and *IRS1*, showed reduced accumulation during infection of cells expressing enzymatically competent myr-Akt relative to the K179M mutant control setting.

Defects in the expression of *UL122 (IE2)* but not *UL123 (IE1)* often relate to the late phase of IE2 expression, which unlike IE1 is sensitive to defects in viral DNA synthesis (28–32). Therefore, we examined if viral DNA synthesis was affected during myr-Akt expression. Indeed, myr-Akt expressing cells showed 6.1-fold reduced levels of in viral DNA at 120 hpi relative to the K179M comparator condition (**Fig. 3C**). The defects in *UL122 (IE2)* mRNA expression and viral DNA synthesis were, as expected, accompanied by decreased protein levels for IE2, as well as the late gene product gB, and the early-late gene product UL44, but not IE1 (**Fig 3C**). Although levels of myr-Akt protein were higher than those seen for the K179M kinase dead control, increased mRNA translation capacity is an expected consequence of constitutive Akt kinase activity [reviewed in (33)]. Overall, these experiments indicate that constitutive Akt activity leads to defective accumulation of viral mRNAs and proteins, and inefficient viral DNA synthesis.

### siRNA knockdown of FOXO3a causes viral replication defects

Akt phosphorylates forkhead box O transcription factors (FOXOs) to prevent their nuclear localization (17, 25, 34), and our previous studies in HCMV latency models found that FOXOs promote reactivation. These observations led us to hypothesize that FOXO transcription factors play roles in lytic cycle progression. If so, the failure of FOXOs to reach the nucleus might contribute to the HCMV lytic cycle defects we observed during expression of constitutively active myr-Akt. To address this possibility, we first turned to short interfering RNA (siRNA) to knock down the expression of either FOXO1 or FOXO3a, two forkhead box O family transcription factors that are expressed in fibroblasts (35). 24 h after transfecting siRNA into fibroblasts, we infected the cells with HCMV strain TB40/E at MOI 1 TCID50/cell, and in parallel experiments evaluated viral replication kinetics, viral DNA synthesis and viral protein expression.

Knockdown of FOXO3a caused a statistically significant roughly 1-log replication defect in the yield of infectious viral particles at 5 days post infection (dpi), as compared to the non-targeting (NT) control siRNA control condition (p=0.0276, **FIG 4A-B)**. The average 5 dpi defect we observed during FOXO3a siRNA treatment was 7.5-fold (range 5.0-fold to 11.2-fold). Silencing of FOXO1 caused a smaller replication defect, on average 2.8-fold at 5 dpi (range 2.2-fold to 3.8-fold), which did not appear to be statistically significant. These viral replication defects were accompanied by inefficient viral DNA synthesis. At 96 hpi, cells silenced for FOXO3a showed a 3-fold reduction in viral DNA accumulation while FOXO1 silenced cells accumulated viral DNA at approximately two-thirds the levels of NT control settings (**FIG 4D-E**).

**Figure 4.**
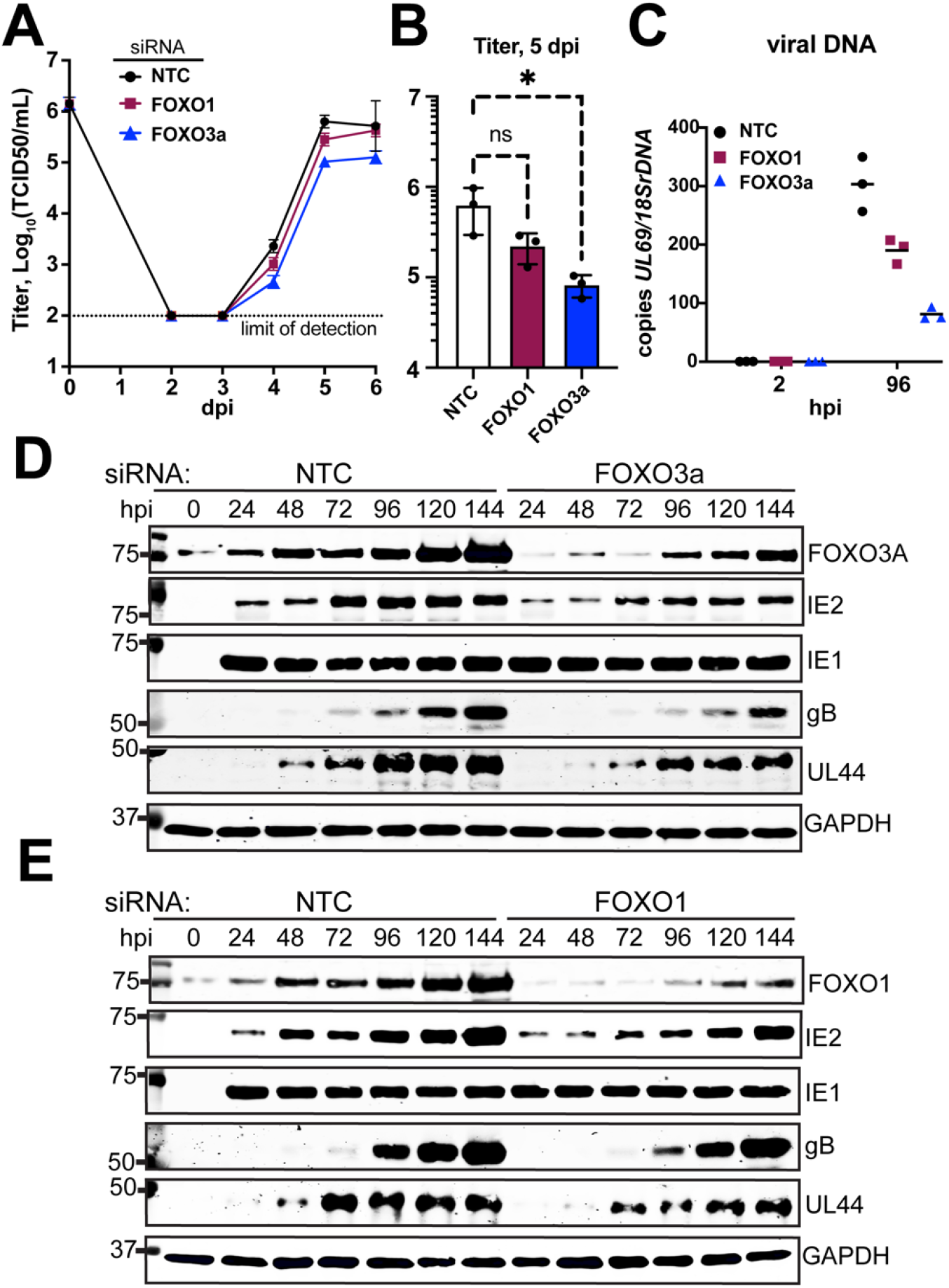
siRNA knockdown of FOXO3a causes viral replication defects. siRNA complexes targeting FOXO1, FOXO3a or both FOXO1 and FOXO3a were reverse transfected into fibroblasts 24 h prior to infection with HCMV strain TB40/E at MOI = 1. **(A)** Supernatants were harvested at the indicated times post infection (days post infection, dpi). **(B) Viral yield at** day 5 post infection for 3 independent biological replicates of the experiment shown in panel A, asterisk indicates p=0.0276 using one-way ANOVA with Dunnet’s post-test to compare means of each siRNA treatment condition to NT control setting. **(C)** Viral DNA copies were enumerated by qPCR using a primer pair specific for HCMV *UL69* and are shown normalized to copies of cellular *18S rDNA* loci. Results are shown for samples collected at 2 h post infection (hpi) and 96 hpi to accurately indicate input levels of viral genomes against replicated viral DNA. **(D-E)** Protein lysates were obtained at the indicated times post infection and analyzed by Western blot to validate knockdown of FOXO3a or FOXO1, respectively, and to assess for effects on viral protein expression. Cellular GAPDH protein was detected as a loading control. NTC: non-targeting control siRNA.

Turning to protein expression, we were intrigued to observe that FOXO3a and FOXO1 protein levels were strongly upregulated during HCMV infection. Normalizing fluorescent detection signals for each FOXO transcription factor to those for GAPDH, allow us to estimate that FOXO1 and FOXO3a were 4.4-fold and 3.0-fold more abundant at 120 hpi, respectively, relative to uninfected cells (**FIG 4D-E, SI Fig S3**). Although the siRNA treatments kept FOXO protein levels below those seen in NT control treated infections, and quantification of secondary antibody fluorophore signals for detection of FOXO1 and FOXO3a at 120 hpi in the siRNA treatment settings indicated that the siRNA treatments reduced their expression by 60% relative to the equivalent time point in the NT control setting (**FIG 4D-E, SI Fig S3**). However, given that knockdown was incomplete, it seems likely that more complete silencing of FOXO expression would lead to greater defects in viral replication and viral DNA synthesis.

In agreement with the viral replication and DNA synthesis defects being greater during silencing of FOXO3a than FOXO1, FOXO3a silenced cells showed poorer expression of glycoprotein B (gB), a late gene (**FIG 4D-4E**). Accumulation of the viral DNA polymerase accessory factor UL44, which exhibits both early and late expression kinetics also showed a more severe decrease relative to the NT control setting in FOXO3a silenced settings. At 120 hpi, GAPDH-normalized detection signals for gB and UL44 in FOXO3a siRNA treated cells were decreased by 54% and 37% relative to NT control infections, respectively. Reduced expression of IE2 was also observed during FOXO3a silencing across all time points. The GAPDH normalized IE2 signal was reduced by 57% relative to NT control at 24 hpi, by 46% at 96 hpi, and by 43% at 120 hpi (**Fig 4D, SI Fig S3**). Although FOXO1 silenced cells likewise showed delayed accumulation of UL44 protein, as well as lower levels of IE2 compared to the NT control setting, these defects were more subtle than what we observed for FOXO3a (**FIG 4E, SI Fig S3**). From these experiments we conclude that silencing of FOXO transcription factors can cause viral DNA synthesis and viral protein expression defects, but that these appear to be more pronounced when FOXO3a is silenced.

### Forced nuclear localization of FOXO3a reverses viral replication defects caused by expression of constitutive AKT

Our siRNA results suggested a more pronounced requirement for FOXO3a than FOXO1 during HCMV lytic infection (**FIG 4**), echoing the role we found for FOXO3a in HCMV reactivation from latency in the THP-1 model and in CD34+ HPCs (19, 20). Therefore, we next sought to interrogate the specific contribution of FOXO3a to the AKT-dependent growth defect we observed during lytic infection. A number of groups have identified FOXO3a regulated cellular genes using a chimeric FOXO3a, dubbed ‘FOXO3a-TM-ER,’ in which FOXO3a is fused to the hormone binding domain of a mutant murine estrogen receptor (ER) carrying an G525R substitution. This mutant form of ER has 1000-fold reduced affinity for estrogen and is instead regulated by 4-hydroxytamoxifen (4-OHT), a synthetic anti-estrogen (36, 37). Further, all three AKT phosphoacceptor sites on FOXO3a are mutated from Ser or Thr to Ala (triple mutant, ‘TM’). Therefore, nuclear localization of the FOXO3a-TM-ER fusion protein is controlled by the addition of 4-OHT, not AKT (**Fig. 5A**). We introduced FOXO3a-TM-ER expression into ARPE-19 retinal pigment epithelial cells that already contained a “tet-on” myr-AKT cassette. (We also attempted to transduce our hTERT-immortalized fibroblasts but failed to obtain healthy cell populations after dual-transduction with both lentivirus vectors [data not shown]).

**Figure 5.**
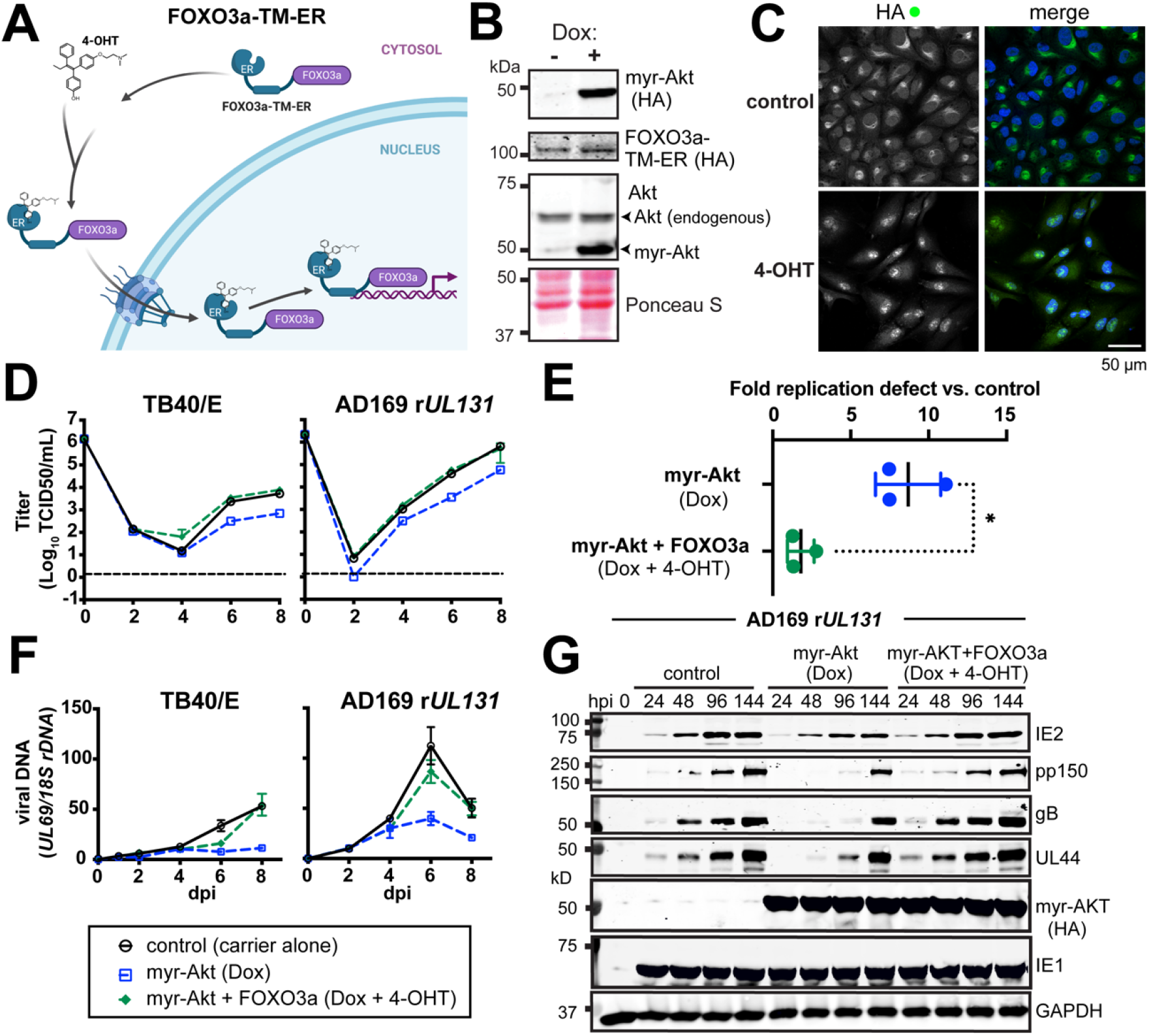
Hormone-regulated chimeric FOXO3a activity reverses AKT-dependent viral replication defects. **(A)** Cartoon depiction (created with Biorender) of chimeric FOXO3a “Triple AKT phosphoacceptor site Mutant” (TM), FOXO3a-TM-ER, in which FOXO3a-TM is fused to a mutant mouse estrogen receptor (ER) in which the anti-estrogen 4-hydroxy-tamoxifen (4-OHT) activates its nuclear localization. Therefore, the FOXO3a-TM-ER protein allows FOXO3a activity to be decoupled from AKT regulation. **(B)** Western blot analysis of ARPE-19 epithelial cells carrying a doxycycline-inducible myr-AKT expression cassette (HA-tagged) and constitutively expressing FOXO3a-TM-ER, also HA-tagged. **(C)** Confocal immunofluorescence microscopy was used to validate 4-OHT induced FOXO3a-TM-ER nuclear localization upon addition of 1 µM 4-OHT for 1 h. HA staining was used to detect the FOXO3a-TM-ER fusion protein; Hoesch 33342 signal is shown to counterstain nuclei the merged image. **(D)** One-step viral replication kinetics at MOI 2 for HCMV strains TB40/E and AD169 repaired for *UL131* (AD169 r*UL131*) in ARPE-19 cells induced for myr-AKT expression (100 ng/mL doxycycline, ‘Dox’) and with or without 4-OHT treatment (1 µM) to induce nuclear localization of FOXO3a-TM-ER, as indicated. **(E)** Results for viral replication yield defect at day 8 post infection from 3 independent biological replicates of strain AD169 r*UL131* in ARPE-19, asterisk indicates p=0.0457 by paired T-test. **(F)** Viral DNA synthesis results for the same time points shown in (**D**), comparing qPCR detected copies of the HCMV *UL69* gene normalized to detection of cellular *18S rDNA* copies. Note, the legend shown atop panel D applies to both D and F; dpi: days post infection. **(G)** Western blot analysis of viral protein expression following AD169 r*UL131* infection (MOI 2 TCID50/mL) in ARPE-19 cells in the presence or absence of Dox and/or 4-OHT treatments, as described above.

We first validated that the FOXO3a-TM-ER ARPE-19 were doxycycline inducible for myr-AKT expression, and that addition of 1 µM 4-OHT resulted in the nuclear localization of the FOXO3a-TM-ER fusion protein (**Fig 5B-C**). Next, we induced myr-Akt expression for 24 h and then infected the cells at MOI 2 TCID50/cell in the presence or absence of 1 µM 4-OHT. In repeated experiments with two different HCMV strains, TB40/E and AD169 r*UL131*, addition of 4-OHT almost entirely rescued the ∼1-log replication defect observed during induction of myr-AKT (**Fig 5D-E, 2A-B**). In three independent biological replicates with AD169 r*UL131*, the fold defect observed at 8 dpi for myr-Akt versus control (carrier alone) conditions was 8.7-fold (range 7.5 - 11.1 fold), but in the presence of 4-OHT the average defect was only 1.8-fold (range 2.8 - 1.3 fold). This difference was statistically significant (p=0.047) by paired, two-tailed t-test (**Fig 5E**). Rescue of viral replication kinetics defects for both viral strains was accompanied by reversal of defects in viral DNA synthesis and viral protein expression that were seen during myr-AKT expression in the absence of 4-OHT (**Fig 5F, 5G**). From these experiments, we conclude that artificial nuclear localization of a FOXO3a is sufficient to reverse the replication defects that occur during expression of constitutive Akt activity.

Collectively, our results argue that FOXO family transcription factors play a critical role in the HCMV lytic replication cycle, and that HCMV relies on Akt inactivation in order to drive FOXOs into the nucleus.

## DISCUSSION

As obligate intracellular parasites, cytolytic viruses routinely re-wire host cell signaling pathways during infection, for instance to upregulate synthesis of macromolecules that support their replication, to evade immune surveillance, and to maintain cell viability while infectious progeny virions are assembled. HCMV is no exception. Although a number of groups have observed that AKT becomes inactivated during HCMV infection (13, 14), it has remained unclear whether cessation of AKT signaling is required for efficient viral replication. However, we have found that constitutively active AKT causes a substantial, ∼10-fold viral replication defect, which is accompanied by substantially reduced late gene expression and defects in viral DNA synthesis (**Fig 2-3**). Our results therefore argue that AKT is necessary for efficient viral replication.

Moreover, data from siRNA knockdown studies (**Fig 4**) and experiments dislocating nuclear localization of FOXO3a from AKT (**Fig 5**), suggest a pivotal role for one AKT substrate in particular, FOXO3a. Indeed, our data show that small molecule (4-OHT)-controlled nuclear localization of FOXO3a rescues virtually all of the viral replication defects caused by constitutive AKT kinase activity. Although we have previously demonstrated that FOXO3a stimulates re-expression of major immediate early genes from intronic promoter elements in the MIE region (19), our new findings argue that FOXO3a in fact plays a broader role in lytic cycle progression, and furthermore that activation of FOXO3a is the major function of AKT inactivation during HCMV infection.

Although our studies here focused on the viral lytic cycle, our findings are also relevant to understanding latency in HCMV and possibly a wide range of herpesviruses. Results from studies of herpes simplex virus 1 (HSV-1), murine gammaherpesvirus-68, human herpesvirus 8 (Kaposi’s sarcoma associated herpesvirus, KSHV), and HCMV collectively underscore a theme in which inhibition of the PI3K-AKT pathway broadly promotes herpesvirus reactivation from latency (14, 38, 39). Further, the KSHV latent protein LANA inhibits FOXO3a activity, which may suggest a role for FOXO3a in KSHV reactivation from latency (40). The latter observation fits well with our previous findings in support of a role for FOXO3a in HCMV reactivation from latency (19, 20). Taken together, our data highlight that PI3K-AKT-FOXO3a may impact dynamic infection states, such as lytic replication versus establishment of latency, as well as progression of the lytic cycle itself, in a broad range of herpesviruses.

Future studies will be needed to elucidate why FOXO3 activity is required during infection beyond transactivation of MIE gene expression. One possibility is that FOXO3a is necessary to upregulate cellular stress response genes, for instance to address the burden of oxidative stress associated with lytic replication. Another, non-mutually exclusive possibility is that HCMV lytic replication genes are directly regulated by FOXO3a, perhaps to regulate viral reactivation from latency in response to extrinsic or intrinsic stressors, much in the same way that cellular genes are regulated by FOXO3a to coordinate stress responses.

## ACKNOWLEDGEMENTS

We are grateful to Thomas E. Shenk (Princeton University) and William J. Britt (University of Alabama, Birmingham), for generously sharing antibody reagents, and to Jacob E. Corn (ETH Zurich), Michael E. Greenberg (Harvard Medical School), Richard A. Roth (Stanford University), William R. Sellers (the Broad Institute), and Didier Trono (École Polytechnique Fédérale de Lausanne), for sharing plasmids via Addgene that were used in this study.

## MATERIALS AND METHODS

### Cells and viruses

Human telomerase reverse transcriptase (hTERT)-immortalized human foreskin fibroblasts, derived from ATCC HFF-1 cells (SCRC-1041), were maintained in Dulbecco’s modified Eagle’s medium supplemented with 5% to 10% newborn calf serum (Millipore Sigma) and antibiotics (complete DMEM) exactly as described previously. HEK-293T cells were purchased from Genhunter Corp. (Nashville, TN). The retinal pigment epithelial cell line ARPE-19 was purchased from ATCC (CRL-2302). All cells were cultured in Dulbecco’s modified Eagle’s medium (DMEM) (Corning 10013CV) supplemented with 25 μg/ml gentamicin, 10 μg/ml ciprofloxacin HCl, and either 5% fetal bovine serum (FBS) (Sigma-Aldrich Cat # F2442 or Gemini Biosciences Foundation B, Cat #900-208) or 5% newborn calf serum (NCS) (newborn calf serum, Gemini Biosciences, #100-504). Studies using doxycycline induction of myr-AKT or myr-AKT K179M, cells were conducted using Opti-MEM medium (Gibco, ThermoFisher) supplemented with antibiotics as above except using tet-approved FBS (Clontech Labs 631106) at 3%.

### Construction of plasmids

Plasmid pLenti-X1-Hygro-mCherry 2A-3xHA FOXO3a TM ER was generated as follows: pLenti-X1 Hygro mCherry RAMP4 (Addgene #118391, a gift of Jacob Corn) was linearized with XcmI and the 10.205 kb fragment was assembled using HiFi Assembly MasterMix (NEB) in a 4-way reaction together with the 2.383 kb NheI/KpnI released FOXO3a-TM-ER fragment from plasmid HA-FOXO3a-TM-ER (Addgene #8353, a gift of Michael Greenberg) and two synthetic dsDNA IDT gBlocks, FOXO3A_ER_gBlock1 (244 bp) and FOXO3A_ER_gBlock2 (248 bp). Full details on sequences of synthetic DNAs generated for this study are provided in **Supplemental Table S1**. (Note: “TM” in FOXO3a-TM indicates “triple mutant” for all three AKT sites, T32A, S253A, S315A). The final lentivirus vector plasmid encodes, under the control of the EF1alpha promoter, mCherry fused to a P2A “self-cleaving” peptide followed by 3 tandem copies of the HA tag (YPYDVPDYA) linked to FOXO3a-TM and then finally fused to the hormone binding domain (amino acids 281-599) of the G525R mutant murine estrogen receptor alpha (41, 42), then terminating with a UGA stop codon.

To construct tet-on plasmids expressing either myr-AKT or myr-AKT-K179M, we digested the plasmid pECE-myrAkt Δ4-129 (Addgene #10841, a gift of Richard A. Roth), with EcoRI-HF and NotI. (Note: Δ4-129 signifies deletion of the AKT pleckstrin homology domain) T4 DNA ligase was then used to insert the released myr-AKT fragment into an agarose gel purified backbone of the all-in-one “tet-on” lentiviral vector plasmid pOUPc_UL148^HA^ (43), which had been opened using EcoRI-HF and NotI to release the *UL148*^*HA*^ insert. This resulted in pOUPc-myrAkt (Δ4-129). A K179M version, pOUPc-K179M myrAkt (Δ4-129) was generated by first double-digesting pECE-myrAkt (Δ4-129) using EcoRI-HF and XbaI to release the 1199 bp fragment containing myr-AKT. This fragment was ligated into the pSP72 vector (Promega) using T4 DNA ligase using the EcoRI and XbaI site on its polylinker, resulting in pSP72-myrAkt (Δ4-129). Then pLNCX myr-Akt_K179M (Addgene #9906, a gift of William R. Sellers) was digested with SmaI + DraIII to release a 289 bp fragment containing the K179M mutation in the context of *AKT1*. The 289 bp fragment was then inserted into a SmaI + DraIII linearized pSP72-myrAkt (Δ4-129) backbone using T4 DNA ligase, yielding pSP72-K179M-myrAkt (Δ4-129). Finally, pSP72 K179 myrAkt (Δ4-129) and pOUPc-myrAkt (Δ4-129) were separately double-digested with EcoRI and BlpI. After agarose gel purification, the 930 bp fragment from pSP72 K179 myrAkt (Δ4-129) containing the K179M mutation was ligated into the 12,341 bp fragment generated after BlpI + EcoRI double-digestion of pOUPc-myrAkt (Δ4-129). All plasmids were confirmed by Sanger sequencing at Genewiz, L.L.C. (Piscataway, NJ).

### Lentivirus vector transduction

All lentiviruses vector particles were generated in HEK-293T cells by co-transfection with plasmids pMD2.g and psPAX2 (Addgene # 12259, #12260, gift of Didier Trono), exactly as described previously (43), except PEI-Max (Polyfect) transfection reagent was used instead of TransIT-293 reagent. To generate APRE-19 cells carrying mCherry_3xHA_FOXO3a_TM_ER, ARPE-19 cells were transduced as described previously (43) and selected in complete DMEM using 50 µg hygromycin B for 3 passages. These cells were then transduced with the “tet-on” lentiviral vector pOUPc-myr-AKT (PH domain, aa 4-129). The cells were further selected with hygromycin (50 µg/mL) and 2 µg/mL puromycin for 3 passages, and then subjected to fluorescence activated cell sorting on a FACSAria III (BD Biosciences) to enrich for mCherry expressing cells. mCherry positive cells were expanded in DMEM containing 10% FBS. During the cell expansion process, selective antibiotics were maintained at 50 µg/ml hygromycin and 2 µg/ml puromycin. The doubly transduced cells were validated by adding 1 µM 4-OHT or ethanol carrier alone control for 1 h and then fixed for HA staining using rabbit anti-HA (Bethyl A190-108A) with goat anti-rabbit Alexa Fluor 488 (Invitrogen, A11008) for secondary detection.

### Western blotting

For Western blot experiments, cells were seeded into 6-well cluster plate at 1×10^6^ cells per well, and treated with doxycycline and/or 4-OHT or carrier controls, and infected with the indicated HCMV strains at MOI 2 TCID50/cell. Cells were lysed at the indicated time points for analysis by Western blot, as described previously (44). Lysis was carried out in 25 mM HEPES (pH 7.5), 400 mM NaCl, 0.1% SDS, 0.5% sodium deoxycholate, 1% NP-40, supplemented with protease inhibitor cocktail (PIC) at 1× (catalog number 5871; Cell Signaling Technologies, Inc., Danvers, MA) and stored at - 80°C. For SDS-PAGE, samples were thawed on ice and then spun at 10,000 × *g* for 30’ at 4°C to remove cell debris. Supernatants were mixed 3:1 with 4× Laemmli buffer (8% sodium dodecyl sulfate [SDS], 0.1% bromophenol blue, 40% glycerol) containing 10% beta mercaptoethanol, denatured at 95°C for 5 min, and resolved on a 10% acrylamide (29:1 acrylamide: bis-acrylamide) SDS-PAGE gel. Proteins on resolved gels were transferred to nitrocellulose membranes (Whatman Protran, 0.45 um pore size). IE2 was detected using mouse monoclonal antibody (mAb) clone 5A8.2 (MAB8140; Millipore, Inc), GAPDH was detected using mouse mAb 60004-1-Ig (Proteintech). All other viral proteins and the HA epitope tag were detected as described previously (44, 45). Details on all antibodies used in this study are provided in **Supplemental Table S2**. All Western blotting results were captured using a LI-COR Odyssey system (LI-COR, Inc., Lincoln, NE). Image Studio 5.25 software (LI-COR) was used to quantify differences in protein expression, with signal from cellular GAPDH being used to normalize signals across lanes, as described elsewhere (43).

### RT-qPCR

For quantification of viral RNA, total RNA was isolated from infected cells using an RNeasy mini kit with an on-column DNase digestion step, as per the manufacturer’s instructions (Qiagen, Inc., Valencia, CA). 1000 ng of total RNA was used as the template to produce oligo(dT)-primed cDNA using qScript cDNA Synthesis kit (Quanta Bioscience, Gaithersburg, MD). Resulting cDNAs and control samples without reverse transcriptase (RT) were diluted 5-fold with water and used as the template in qPCRs to quantify the abundance of the transcripts of the indicated viral genes and of cellular *GAPDH* (glyceraldehyde-3-phosphate dehydrogenase) transcripts, which were used for normalization. cDNA was then used in standard SYBR-green real-time PCR using NEB Luna Master Mix (M3003E) to measure gene expression using the indicated primer pairs (**Supplemental Table S1**), as described previously (43). Reaction conditions were 95°C for 15 s and 60°C for 1 min, repeated for 40 cycles, with a 10-min hot start at 95°C. Relative mRNA levels were quantified using the 2^ΔCt^ method, standardizing to RT-qPCR results for *GAPDH*. Control samples in which reverse transcriptase was intentionally left out during the cDNA synthesis step uniformly produced qPCR results indistinguishable from the results of RT-qPCR on RNA samples from mock-infected cells (not shown).

### siRNA reverse transfection

siRNAs were reverse transfected into hTERT immortalized HFF cells using Lipofectamine RNAiMAX reagent (Thermo Fisher) according to the manufacturer’s instructions. Briefly, two mixes were prepared separately. Mix 1 was prepared by adding 30 pmol of siRNA to 150 µl non-supplemented Opti-MEM medium (Thermo Fisher) and then gently mixing. Mix 2 was prepared by adding 9 µl of RNAiMAX reagent to 150 µl non-supplemented Opti-MEM. Mixes 1 and 2 were then combined, immediately transferred to an empty well of a 6-well plate, and incubated at room temperature for 5 min. Approximately one-million cells were then added to the well in 2.0 ml of DMEM containing 5% NCS (10 nM siRNA final). 24 h later, the cells were infected at an MOI of 1 TCID50 per cell with HCMV TB40/E strain. 2 h after infection, cells were washed twice in PBS (137 mM NaCl, 2.7 mM KCl, 10 mM Na_2_HPO_4_, and 1.8 mM KH_2_PO_4_, pH 7.4), allowing the PBS to sit on the cells for 5 min each time, and then fresh complete DMEM (5% NCS) was added. At the indicated time points, supernatants were collected for viral titer determination, siRNA used in this study were: Dharmacon Non Targeting Control Pool #2 (Cat # D-001206-14-05), SignalSilence® FoxO1 siRNA I (Cell Signaling Tech, Cat# 6242), SignalSilence® FoxO1 siRNA I (Cell Signaling TechCat # 6302). RNA was extracted from HFF at indicated time points using the RNeasy Mini Kit (Qiagen, Hilden, Germany).

### Induction of FOXO-TM-ER fusion protein nuclear localization

For the experiment in Fig 5, ARPE–19 cells were seeded at 1×10^5^ cells per well in 24-well cluster plates. Where indicated, cells were treated with doxycycline (100 ng/ml) for 24 h, and then treated with (Z)-4-Hydroxytamoxifen (4-OHT, Sigma-Aldrich Cat# 508225) at a final concentration of 1 µM for 1 h, to induce myr-AKT or nuclear localization of FOXO3a-TM-ER, respectively. Each drug was applied from a 1000× stock solution (in water for doxycycline, and in ethanol for 4-OHT), therefore, carrier alone control wells were treated with the same volume of absolute ethanol and/or water (0.1% final). Next, viruses TB40/E or AD169 *rUL131* were applied at MOI=2 (2 TCID50 per well), as indicated. For viral replication kinetics studies, infected cell supernatants were collected at the indicated time points and stored at -80°C until determination of titer by TCID50 assay using the Reed Muench method, as described elsewhere (45).

### Viral DNA synthesis assays

Total DNA was isolated from infected cells using the DNeasy Blood and Tissue mini kit (Qiagen, Cat# 69506). Briefly, cells were seeded into 24-well plates, as described above for replication kinetics studies, and at the indicated times postinfection, culture media was removed, and cells were washed three times with PBS, allowing PBS to sit on cells for 5 min per wash. After the PBS wash was aspirated, 200 µL Qiagen buffer AL was added to well, followed by 200 µL PBS and 20 µL of the proteinase K solution included with the kit. Cells were then scraped from the wells using a 1000 µL capacity pipette tip to mix them into the lysis buffer solution. Lysates were then transferred to 1.5 mL microcentrifuge tubes and processed as per manufacturer instructions to isolate total DNA. Absolute quantification of viral DNA copies with specific primers targeting the viral *UL69* gene was performed using quantitative PCR (qPCR) alongside duplicate standard curves of serial ten-fold dilutions of TB40/E BAC DNA from 10^7^ to 10^2^ copies per well for determination of viral copy number, using the same qPCR reagents described above (see RT-qPCR). Meanwhile, a separate standard curve was generated using a plasmid containing a fragment of the human *18S* rDNA gene, from 10^7^ to 10^2^ copies per well, to allow absolute quantification of *18S* rDNA copies. Viral *UL69* copies were then normalized across samples to cellular *18S* rDNA copies (primer details shown in **Supplemental Table S1)**.

### Confocal microscopy and immunofluorescence

Confocal immunofluorescence was carried out essentially as described previously (44). For the experiment shown in Fig 1, hTERT-immortalized human foreskin fibroblasts were seeded on 12-mm, circular, number-1 thickness microscope cover glass (200121; Azer Scientific, Morgantown, PA) and infected with HCMV strain TB40/E at MOI=1 at 37°C in a humidified 5% CO_2_ incubator. 2 h later, culture medium was removed, cells were washed twice in PBS, and then incubated in fresh complete medium containing 5% NCS. Cells were then fixed at the indicated times post infection by removing culture media, washing with PBS and then applying a solution of 4% paraformaldehyde (PFA) in PBS for a minimum of 20 min at room temperature. After fixation, PFA was removed and cells were washed three times in PBS, allowing 5 min for each wash. Cells were then permeabilized for 10 min using 0.1% Triton X-100 (in PBS) at room temperature, and then washed an additional three times in PBS. After washing, cells were blocked at 37°C for 45 min in a solution of 5% normal goat serum (Rockland Immunochemicals) in PBS, then washed another three times in PBS, and then additionally blocked using 1% human BD Fc block (BD Biosciences) in PBS at 37°C for an addition 45 min. Cells were then washed against three times in PBS, for 5 min per wash, and then incubated with antibodies specific for IE1 and FOXO3a (Cell Signaling Technology) at 4°C overnight (antibody details provided in **Supplemental Table S2**). The next day, cells were washed in PBST (PBS containing 0.1% Tween-20). After three washes in PBST, cells were incubated with AlexaFluor 488 conjugated goat anti-rabbit and AlexaFluor 405 conjugated goat anti-mouse secondary antibodies (Invitrogen). After secondary antibody incubation, cells were washed three times with PBST, 5 min per wash. Next, Prolong™ anti-fade mounting medium without DAPI (ThermoFisher) was applied to a slide and the cover slip was inverted onto the mounting media and sealed with nail polish around the edge of the coverslip. Confocal imaging was obtained on a Leica TCS SP5 confocal microscope using a 63× oil immersion lens (Leica Microsystems). In Fig 5C, the same staining procedure was used except the ARPE-19 cells were not infected, and anti-HA antibody (Bethyl) and Alexa488 conjugated goat anti rabbit secondary antibody were used to detect FOXO3a-TM-ER by virtue of its 3× HA epitope tag. Finally, nuclei were counterstained using a ten-minute incubation in PBS containing 10 µg/ml Hoechst 33342 (Millipore Sigma, Cat # 14533). Cells were then washed three times in PBS and mounted in Prolong™ anti-fade mounting medium without DAPI.

---

**SI Figure 1.**
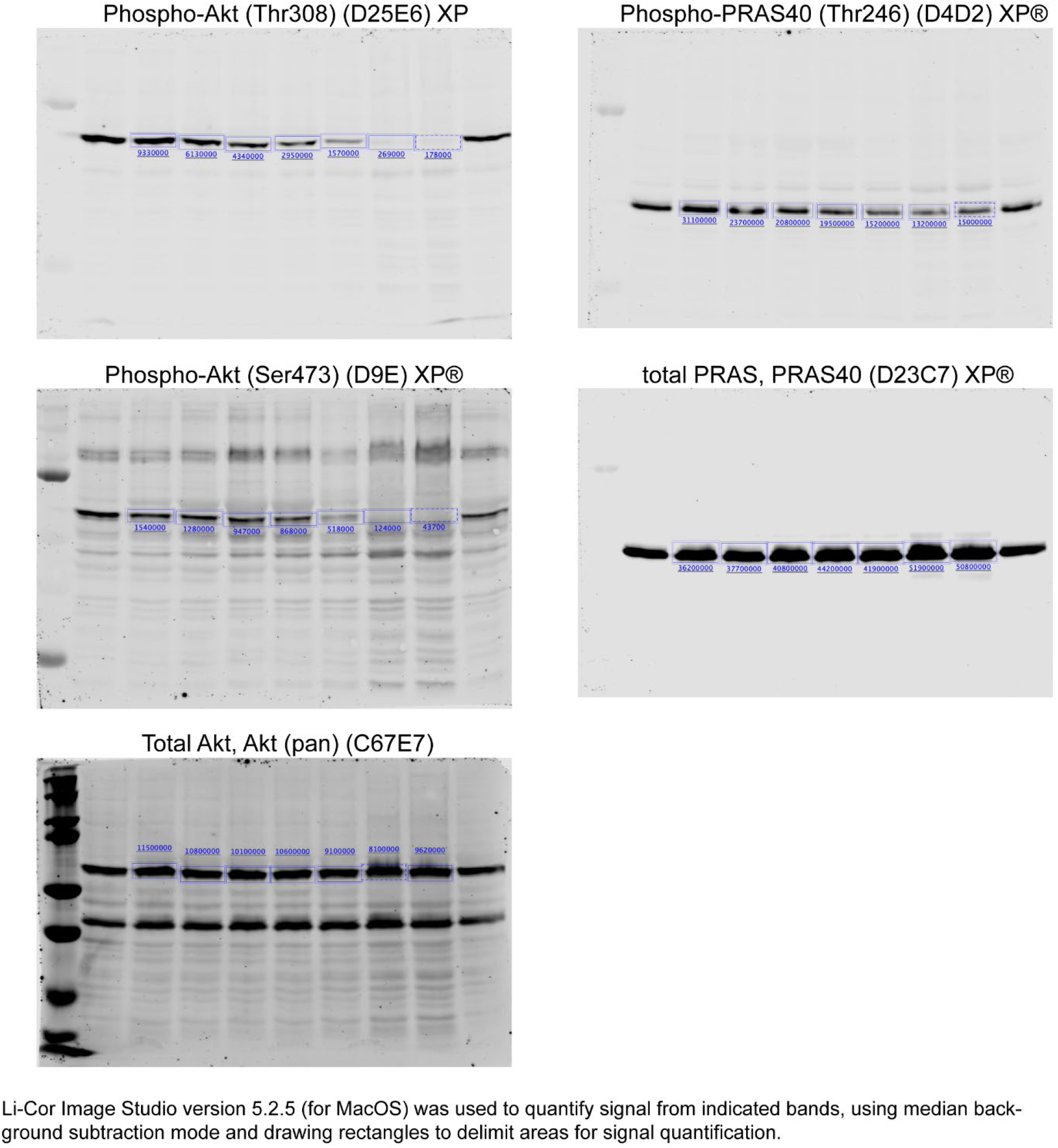
Li-Cor quantification of Western blot results from Figure 1.

**SI Figure 2.**
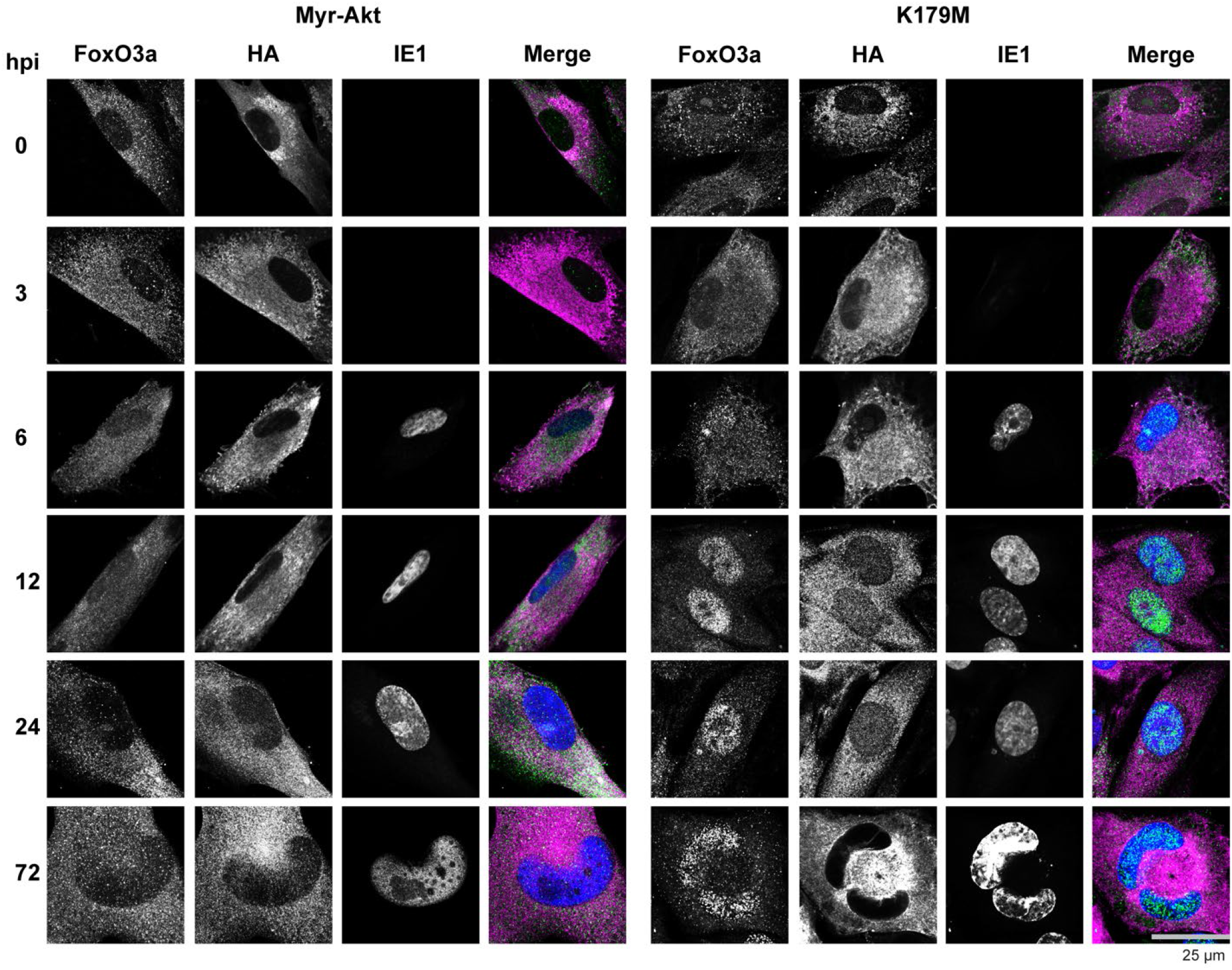
Full image time course series of the results shown in Figure 2C.

**SI Figure 3.**
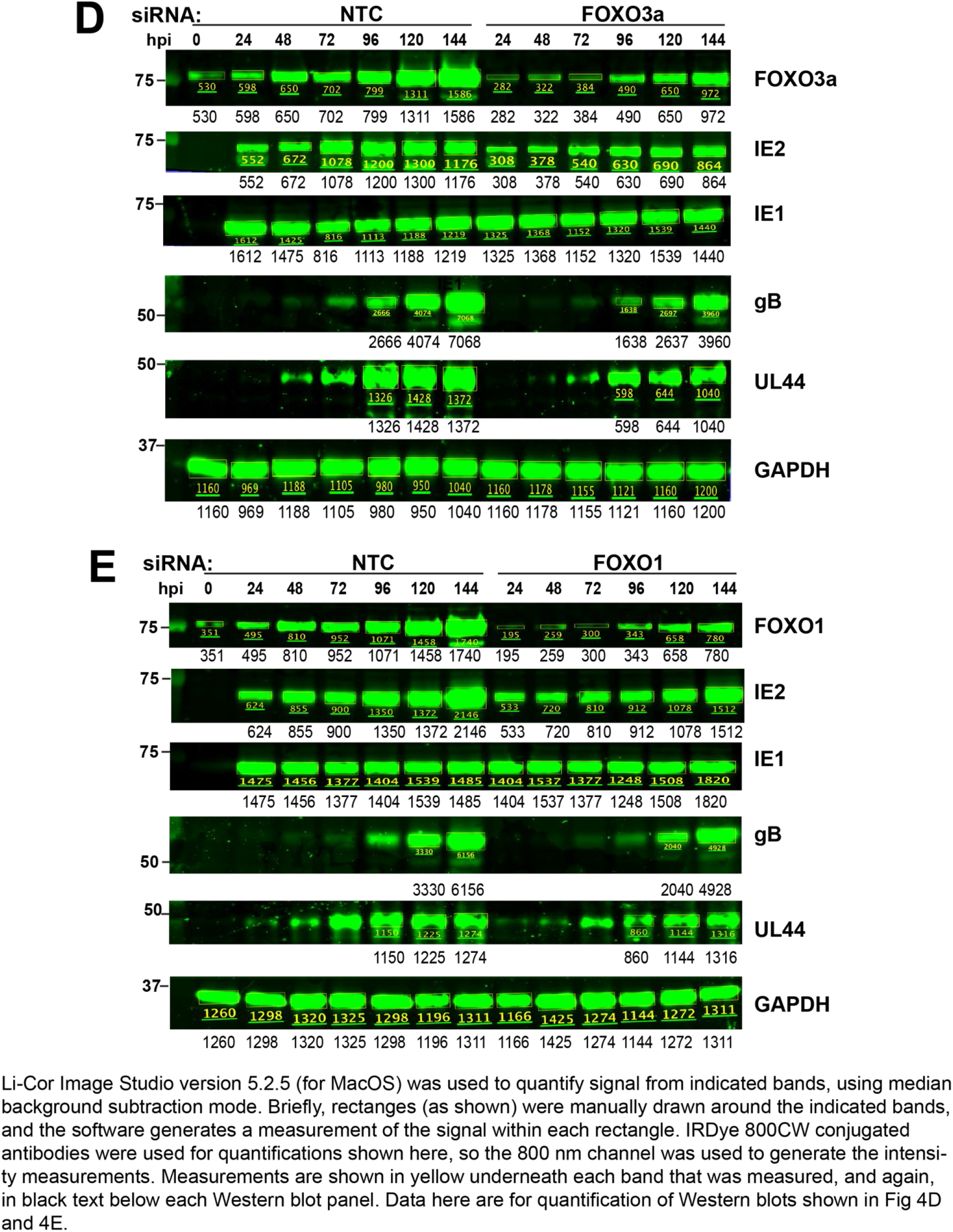
Li-Cor quantification of Western blot results from Figure 4D-4E.

**Supplemental Table S1.**
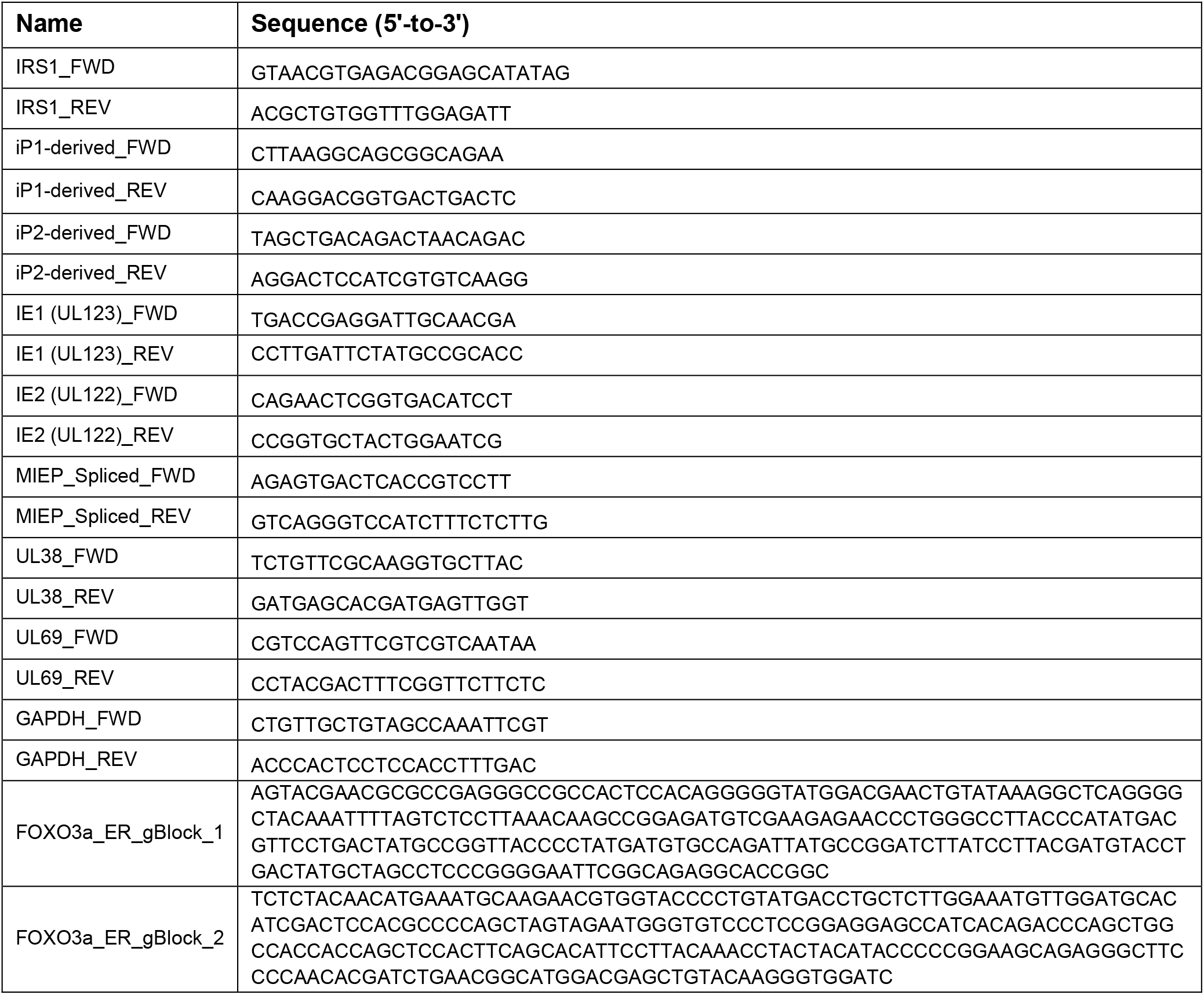
qPCR primers and synthetic custom deoxynucleotide sequences used in this study.

**SUPPLEMENTAL TABLE S2.** Antibodies used in this study

| Primary Antibody | Host species | Isotype | Clone | Source | Catalog No. | Dilution used |
| --- | --- | --- | --- | --- | --- | --- |
| HA epitope | Chicken | IgY |  | Bethyl | A190-106A | 1:200 |
| HCMV major immediate-early (IE) protein IE1-72kD | Mouse | Unknown IgG | 1B12 | Thomas Shenk | n/a | 1: 50 |
| HCMV major immediate-early (IE) protein IE2-86kD | Mouse | IgG1 | 5A8.2 | Millipore, Inc. | MAB8410 | 1:1000 |
| HA epitope | Rabbit | pAb |  | Bethyl, Inc. | A190-108A | 1:1000 |
| FoxO3a (D19A7) | Rabbit | mAb |  | Cell Signaling Technology | 12829S | 1:200 |
| FoxO1 (C29H4) | Rabbit | mAb |  | Cell Signaling Technology | 2880S | 1:1000 |
| Akt (pan) (C67E7) | Rabbit | mAb |  | Cell Signaling Technology | 4691S | 1:1000 |
| PRAS40 (D23C7) XP® | Rabbit | mAb |  | Cell Signaling Technology | 2691S | 1:1000 |
| Phospho-Akt (Thr308) (D25E6) XP® | Rabbit | mAb |  | Cell Signaling Technology | 13038S | 1:1000 |
| Phospho-Akt (Ser473) (D9E) XP® | Rabbit | mAb |  | Cell Signaling Technology | 4060S | 1:1000 |
| Phospho-PRAS40 (Thr246) (D4D2) XP® | Rabbit | mAb |  | Cell Signaling Technology | 13175S | 1:1000 |
| gB | Mouse | mAb | 27-180 | William J. Britt | n/a | 1:1000 |
| UL44 | Mouse | IgG1 | 10D8 | Virusys, Inc. | CA006-100 | 1:1000 |
| GAPDH | Mouse | IgG2b |  | Proteintech | 60004-1-Ig | 1:20,000 |
| Secondary Antibody | Host species | Isotype |  | Vendor | Catalog No. | Dilution used |
| Alexa Fluor 488 anti-Mouse IgG (H+L), cross adsorbed | Goat | IgG |  | ThermoFisher / Invitrogen | A11001 | 1:1000 |
| Alexa Fluor 488 anti-Rabbit IgG (H+L) Cross-Adsorbed | Goat | IgG |  | ThermoFisher / Invitrogen | A11008 | 1:1000 |
| Alexa Fluor 594 anti-Rabbit IgG (H+L) Cross-Adsorbed | Goat | IgG |  | ThermoFisher / Invitrogen | A11012 | 1:1000 |
| Alexa Fluor 594 anti-Mouse IgG (H+L) Cross-Adsorbed | Goat | IgG |  | ThermoFisher / Invitrogen | A11005 | 1:1000 |
| Alexa Fluor 647 anti-Chicken IgY (H+L) | Goat | IgG |  | ThermoFisher / Invitrogen | A11039 | 1:1000 |

